# An encoding model of temporal processing in human visual cortex

**DOI:** 10.1101/108985

**Authors:** Anthony Stigliani, Brianna Jeska, Kalanit Grill-Spector

**Affiliations:** Department of Psychology, Stanford University, Stanford, CA 94305, USA; Stanford Neurosciences Institute, Stanford University, Stanford, CA 94305, USA

**Keywords:** fMRI, V1, V4, MT, extrastriate cortex, temporal channels, transient, sustained

## Abstract

How is temporal information processed in human visual cortex? There is intense debate as to how sustained and transient temporal channels contribute to visual processing beyond V1. Using fMRI, we measured cortical responses to time-varying stimuli, then implemented a novel 2 temporal-channel encoding model to estimate the contributions of each channel. The model predicts cortical responses to time-varying stimuli from milliseconds to seconds and reveals that (i) lateral occipito-temporal regions and peripheral early visual cortex are dominated by transient responses, and (ii) ventral occipito-temporal regions and central early visual cortex are not only driven by both channels, but that transient responses exceed the sustained. These findings resolve an outstanding debate and elucidate temporal processing in human visual cortex. Importantly, this approach has vast implications because it can be applied with fMRI to decipher neural computations in millisecond resolution in any part of the brain.

---

How does the visual system process the temporal aspects of the visual input? In the retina^1^ and LGN^2-4^ temporal processing is thought to be mediated predominately by a magnocellular (M) pathway distinguished by its large transient responses^3,4^ and a parvocelluar (P) pathway which has larger sustained responses than the M pathway^3,4^ (in addition to a smaller koniocelluar^5^ pathway). While M and P pathways remain segregated up to striate cortex (V1), there is intense debate as to how these pathways contribute to visual processing in extrastriate cortex. The prevailing view suggests that the dorsal stream, particularly MT, is M dominated^6-8^, and the ventral stream, particularly V4, is P dominated^9-11^. However, an opposing view suggests that these pathways are not segregated in extrastriate cortex^5, 8^ as there is evidence for M and P contributions to both V4^5, 9^ and MT^12, 13^.

Since M and P pathways are associated with transient and sustained responses, respectively, these theories make predictions regarding temporal processing in human visual cortex. The prevailing view predicts that human MT complex (hMT+) will have large transient but small sustained responses, and conversely human V4 (hV4) will have large sustained but small transient responses. However, the opposing view predicts substantial transient and sustained responses in both hMT+ and hV4. While these predictions are derived from studies of the macaque brain, whether the same predictions can be made to the human brain is uncertain because the organization of human visual cortex differs from the macaque in three notable ways: (1) V4 and MT neighbor in the macaque brain, but hV4 and hMT+ are separated by ~3 cm in the human brain, (2) whether macaque V4 and hV4 are homologous is subject to debate^14-17^, and (3) the human brain contains several additional visual regions neighboring hV4 and hMT+ that are not found in the macaque (VO-1/VO-2 and LO-1/LO-2, respectively). Thus, generating a complete model of temporal processing in human visual cortex necessitates measurements in humans.

Understanding temporal processing in human visual cortex has seen little progress for two main reasons. First, the temporal resolution of fMRI is in the order of seconds^18^, an order of magnitude longer than the timescale of neural processing, which is in the tens to hundreds of milliseconds range. Second, while fMRI responses are largely linear for long stimulus presentations^19, 20^, they exhibit marked nonlinearities for short and transient stimuli^19-24^. Since the standard linear model for fMRI^19, 25^ is inadequate for modeling responses to such stimuli and fMRI is slow, the temporal processing characteristics of human visual cortex remain elusive.

If the observed nonlinearities are of neural (rather than BOLD) origin, a new encoding approach applied to fMRI^26-29^, which uses computational models to predict neural responses (even if they are nonlinear), could surmount these issues. Different than the standard method, which predicts fMRI signals directly from the stimulus, the encoding approach first models neural responses to the stimulus, then from the predicted neural responses calculates fMRI responses. The encoding approach^26-29^ has been influential for two reasons: (i) it provided an important insight that accurately modeling neural responses at a sub-voxel resolution better predicts fMRI responses at the voxel resolution, and (ii) it advanced understanding of neural mechanisms by building explicit, quantitative models of neural computations.

Here we sought to leverage the encoding approach to characterize temporal processing in human visual cortex. Thus, we built a temporal encoding model of neural responses to time-varying visual stimuli in millisecond resolution and used this model to predict fMRI responses in second resolution. The model is based on estimation of the transient and sustained channels’ impulse response functions from measurements in macaque V1^30-32^ and psychophysics in humans^33-36^. To determine temporal processing in human visual cortex we implemented three experiments aimed to measure fMRI responses to time-varying visual stimuli that were either sustained (1 continuous image per trial, durations ranging from 2 s to 30 s; **Fig. 1**, *Experiment 1*), transient (30 flashed, 33 ms long, images per trial, inter-stimulus intervals ranging from 33–967 ms; **Fig. 1**, *Experiment 2*), or contained both transient and sustained components (30 continuous images per trial, durations ranging from 67–1000 ms per image; **Fig. 1**, *Experiment 3*). We first determined if millisecond temporal variations in visual stimuli generate substantial modulations of fMRI responses in visual cortex. Then we used Experiments 1 and 2 to estimate the 2 temporal-channel encoding model parameters and tested how well the model predicts fMRI responses to stimuli that vary in their temporal properties from milliseconds to seconds in new data from Experiment 3. Once we established the model’s validity, we derived the contributions of sustained and transient channels to neural responses across striate and extrastriate visual cortex to test the competing experimental hypotheses.

**Figure 1:**
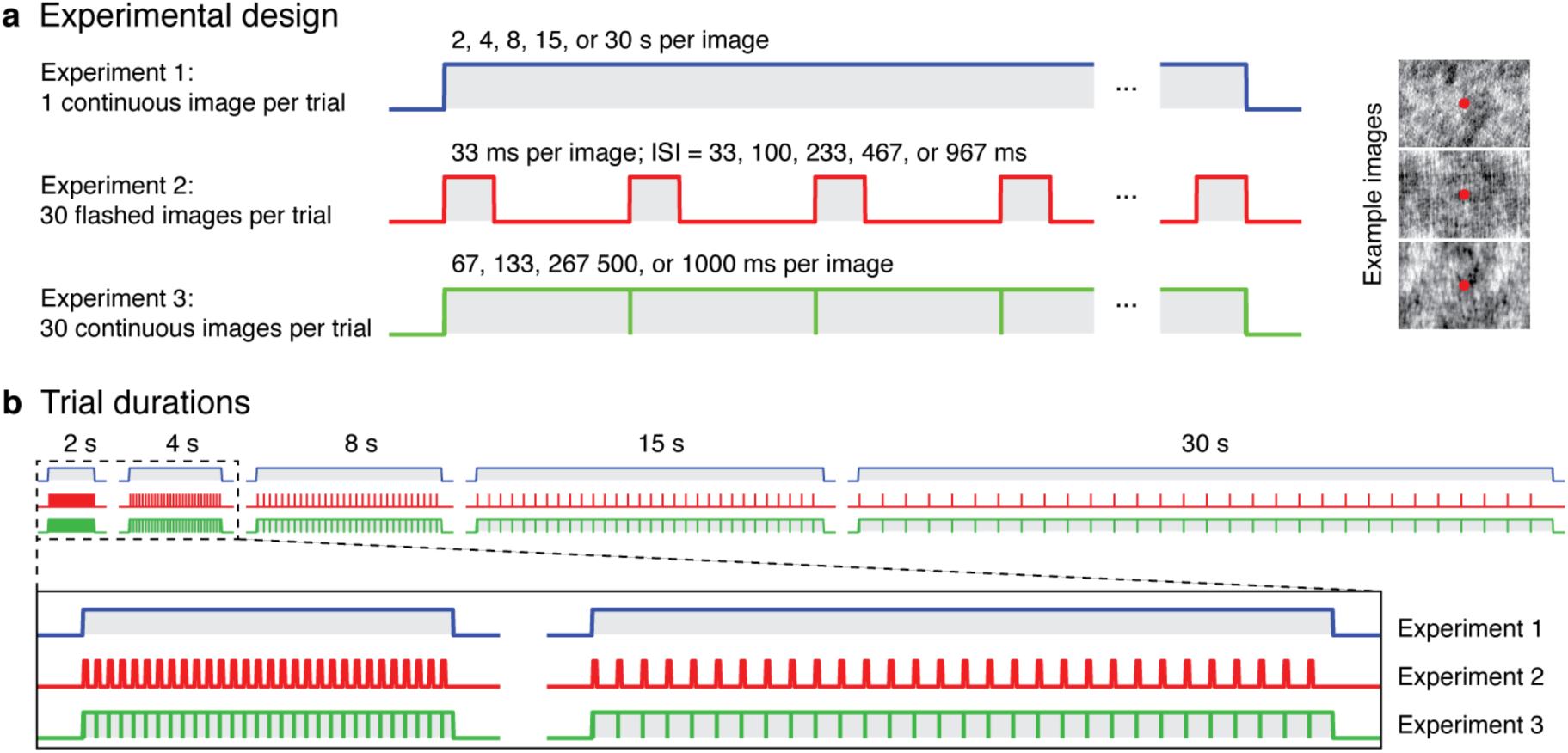
Measuring brain responses to combinations of sustained and transient visual stimuli. (**a**) Participants fixated centrally and viewed phase-scrambled gray-level images that were presented in trials of different durations, interleaved with 12 s periods of a blank screen. The same fixation task (detecting change in fixation color) was used in all three experiments. *Experiment 1:* a single phase scrambled image was shown for the duration of a trial. *Experiment 2:* 30 briefly presented images (33 ms each), each followed by a blank screen, were presented in each trial. As the trial duration lengthens, the gap between images increases, causing the fraction of the trial containing visual stimulation to decrease. *Experiment 3:* 30 continuous images (with no gaps between consecutive stimuli) were presented in each trial. As the block duration lengthens, the duration of each image progressively increases. (**b**) The same trial durations (2, 4, 8, 15 or 30 s) were utilized across all three experiments while the rate and duration of visual presentation varied between experiments. Corresponding trials in Experiments 1 and 3 have the same overall duration of stimulation but different numbers of stimuli, whereas trials in Experiments 2 and 3 have the same number of stimuli but different durations of stimulation. *Top:* stimulation durations for example trial in each experiment; *Bottom:* zoom on the 2 s and 4 s trials.

## RESULTS

### Do Millisecond Temporal Variations in the Visual Stimulus Modulate V1 Responses

To test the feasibility of this approach, we first examined V1 responses during the three experiments. Predicted fMRI responses from the standard model depend only on the type and duration of stimuli. Thus, the standard model predicts longer responses for longer trials and identical responses in Experiments 1 and 3 (**Fig. 2a**, *blue and green*), which use the same visual stimuli and trial durations and just vary by the number of images per trial (1 vs. 30, respectively). Furthermore, the model predicts that the amplitude of responses in Experiments 1 and 3 will increase from 2–8 s trials and will remain largely the same for longer trials. While the standard model predicts the same response durations in Experiment 2, it predicts substantially lower response amplitudes in Experiment 2 than Experiments 1 and 3 because the transient visual stimuli are presented for only a fraction of each trial. Furthermore, the model predicts a progressive decrease in response amplitude during Experiment 2 from 2–30 s trials as the fraction of the trial in which visual stimuli are presented decreases (from 1/2 to 1/30 of the trial, **Fig. 2a**, *red*).

**Figure 2:**
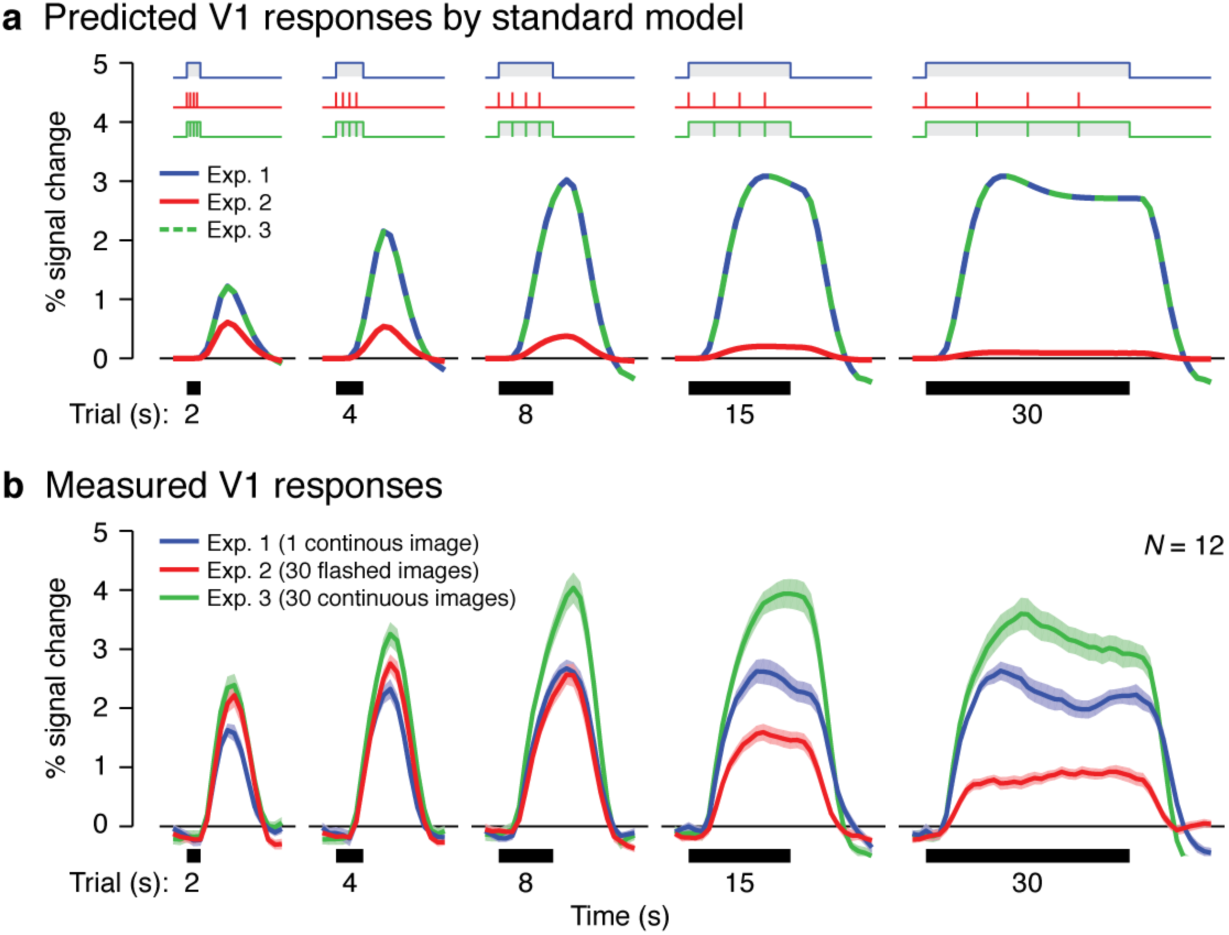
V1 responses to transient stimuli differ from the predictions of the standard model. (**a**) The standard model predicts the same response in trials of the same duration across Experiment 1 (*blue*) and Experiment 3 (*green*), since both present stimuli continuously for the same total duration in each trial. However, responses in Experiment 2 (*red*) are predicted to be much lower because stimuli are spaced apart and are only presented for a fraction of each trial duration. (**b**) The mean V1 response in Experiments 1–3 averaged across 12 participants. Each curve is data from a different experiment (see legend). Shaded regions around the curves indicate ± 1 standard error of the mean (SEM) across 12 participants. In both (a) and (b) the onsets and lengths of the trials are illustrated as thick black bars below each graph, and curves extend 2 s before the onset and 12 s after the offset each trial.

While V1 responses to sustained visual stimulation in Experiment 1 largely followed the predictions of the standard model (**Fig. 2a**, *blue*), responses in Experiments 2 and 3 deviated from the standard model’s predictions. First, responses in Experiment 3 (**Fig. 2b**, *green*) were higher than responses in Experiment 1 for all trial durations. Second, responses to transient stimuli in Experiment 2 (**Fig. 2b**, *red*) were substantially higher than predicted by the standard model. In fact, V1 responses during 2–8 s trials of Experiment 2 were equal or higher than those of Experiment 1, even though the cumulative duration of stimulation across images in Experiment 2 was a fraction of the duration of stimulation in Experiment 1. Third, different than the predictions of the standard model, response amplitudes in Experiment 2 did not systematically decline with trial duration but instead peaked for the 4 s trials.

These data demonstrate that (i) varying the temporal characteristics of visual presentation in the millisecond range has profound effects on V1 fMRI responses, and (ii) the standard model is inadequate in predicting measured fMRI responses for these stimuli, in agreement with prior data. Furthermore, the higher responses in Experiment 3 (which has both sustained and transient visual stimulation) compared to Experiments 1 and 2 (which have either sustained or transient stimuli, respectively) suggest that both transient and sustained components of the visual input contribute to the fMRI signals, consistent with our hypothesis.

### An Encoding Model for Temporal Processing in Visual Cortex

To accurately predict fMRI responses in all three experiments, we built a temporal encoding model of neural responses in millisecond resolution and used this model to predict fMRI responses in second resolution (**Fig. 3**). Our model consists of 2 neural temporal channels, each of which can be characterized by a linear systems approach using a temporal impulse response function^30-32, 34-36^ (IRF). The sustained channel is characterized by a monophasic IRF (**Fig. 3b**, *blue channel IRF*) peaking at around 40 ms and lasting 100–150 ms; convolving this channel with a visual stimulus will produce a sustained neural response for the duration of the stimulus. The transient channel is characterized by a biphasic IRF, akin to a derivative function, with the positive part peaking at around 35 ms and the negative part peaking at around 70 ms (**Fig. 3b**, *red channel IRF*). A squaring nonlinearity is added, as both stimulus onset and offset lead to increased neural firing and consequently increased metabolic demands^36, 37^. Thus, convolving the visual stimulus with this transient IRF will produce a positive neural response when there is an onset or offset of the visual stimulus but zero response in between when the stimulus is presented for durations longer than the duration of the IRF. The predicted fMRI response is generated by convolving the output of each neural channel with the HRF and summing the responses of the two temporal channels (**Fig. 3c**).

**Figure 3:**
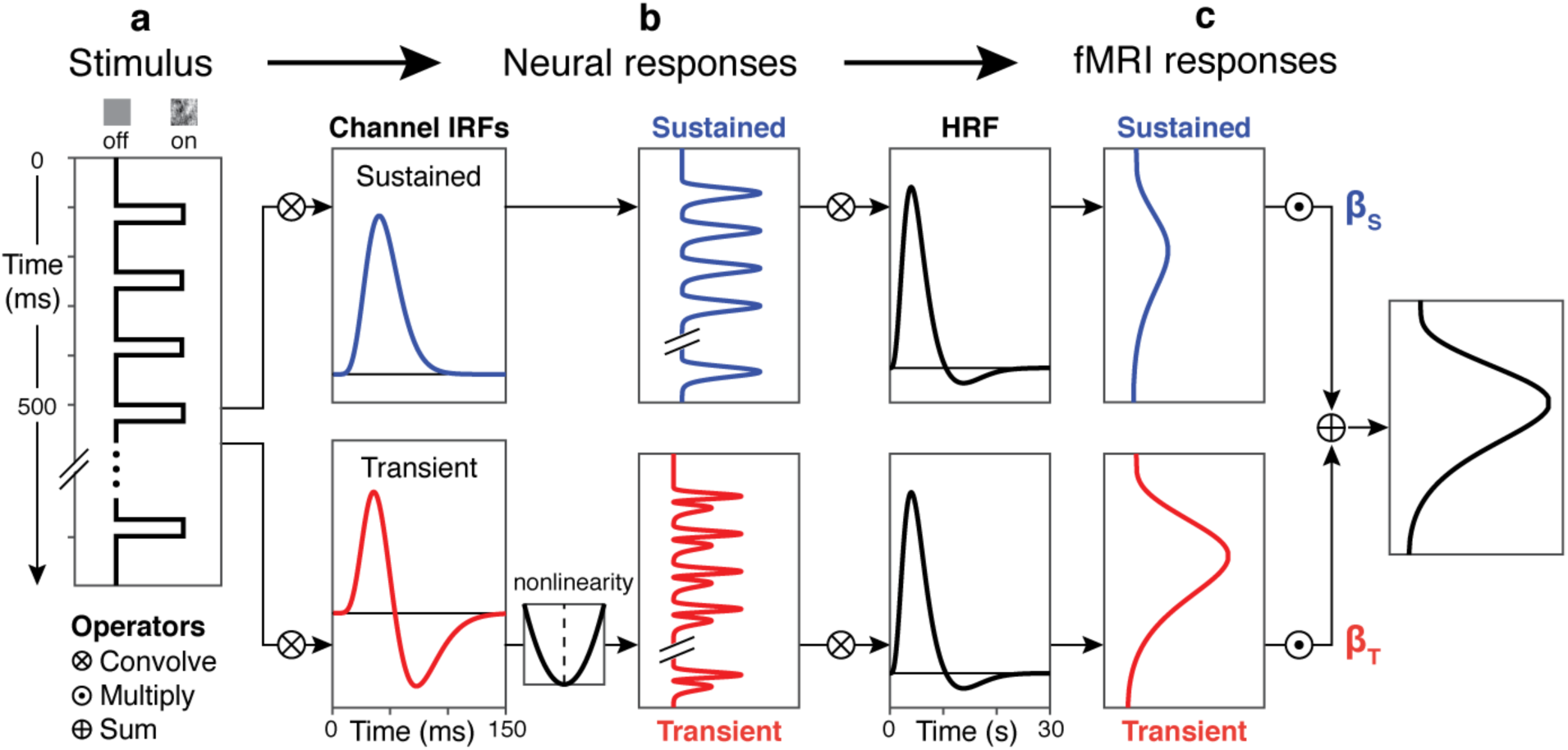
The 2 temporal-channel model. (**a**) Transitions between stimulus and baseline screens are coded as a step function representing when a stimulus was on vs. off with millisecond temporal resolution. In the example illustrated here, each stimulus is presented for 33 ms and followed by a 100 ms blank screen. (**b**) Separate neural responses for the sustained (*blue*) and transient (*red*) channels are modeled by convolving the stimulus vector with impulse response functions (IRFs) for the sustained and transient channels, respectively, estimated from human psychophysics. A squaring nonlinearity is applied in the transient channel to rectify offset deflections (see Online Methods). (**c**) Sustained and transient fMRI response predictors are generated by convolving each channel’s neural responses with the hemodynamic response function (HRF) and down-sampling to match the temporal acquisition rate of fMRI data. The total fMRI response is the sum of the weighted sustained and transient fMRI predictors for each channel. To estimate the contributions (*β* weights) of the sustained (*β_S_*) and transient (*β_T_*) channels in V1, we fit the 2 temporal-channel model across data concatenated across Experiments 1 and 2.

Our procedure for testing the 2 temporal-channel encoding model had two stages. First, we estimated the contributions of the two temporal channels to fMRI signals using concatenated data from Experiments 1 and 2 that were designed to largely drive sustained or transient channels, respectively. Second, we cross-validated the model by testing how well it predicted data from a third experiment that had both transient and sustained visual stimulation. As a benchmark, we compared the performance of the 2 temporal-channel model with the standard model.

### Does a 2 Temporal-Channel Model Explain V1 fMRI Responses to Time-Varying Stimuli?

Comparing the predictions of the 2 temporal-channel model to V1 responses reveals three findings. First, the 2 temporal-channel model containing one sustained predictor (weighted by *β_S_*) and one transient predictor (weighted by *β_T_*) generated fMRI signals that tracked both the duration and amplitude of V1 responses in Experiments 1 and 2 (**Fig. 4a**, compare model prediction, *black,* to measured V1 data, *gray*). Consistent with our predictions, the sustained channel accounted for the majority of responses in Experiment 1 (**Fig. 4a**, *top row, blue*), while the transient channel contributed the bulk of the response in Experiment 2 (**Fig. 4a**, *bottom row, red*).

**Figure 4:**
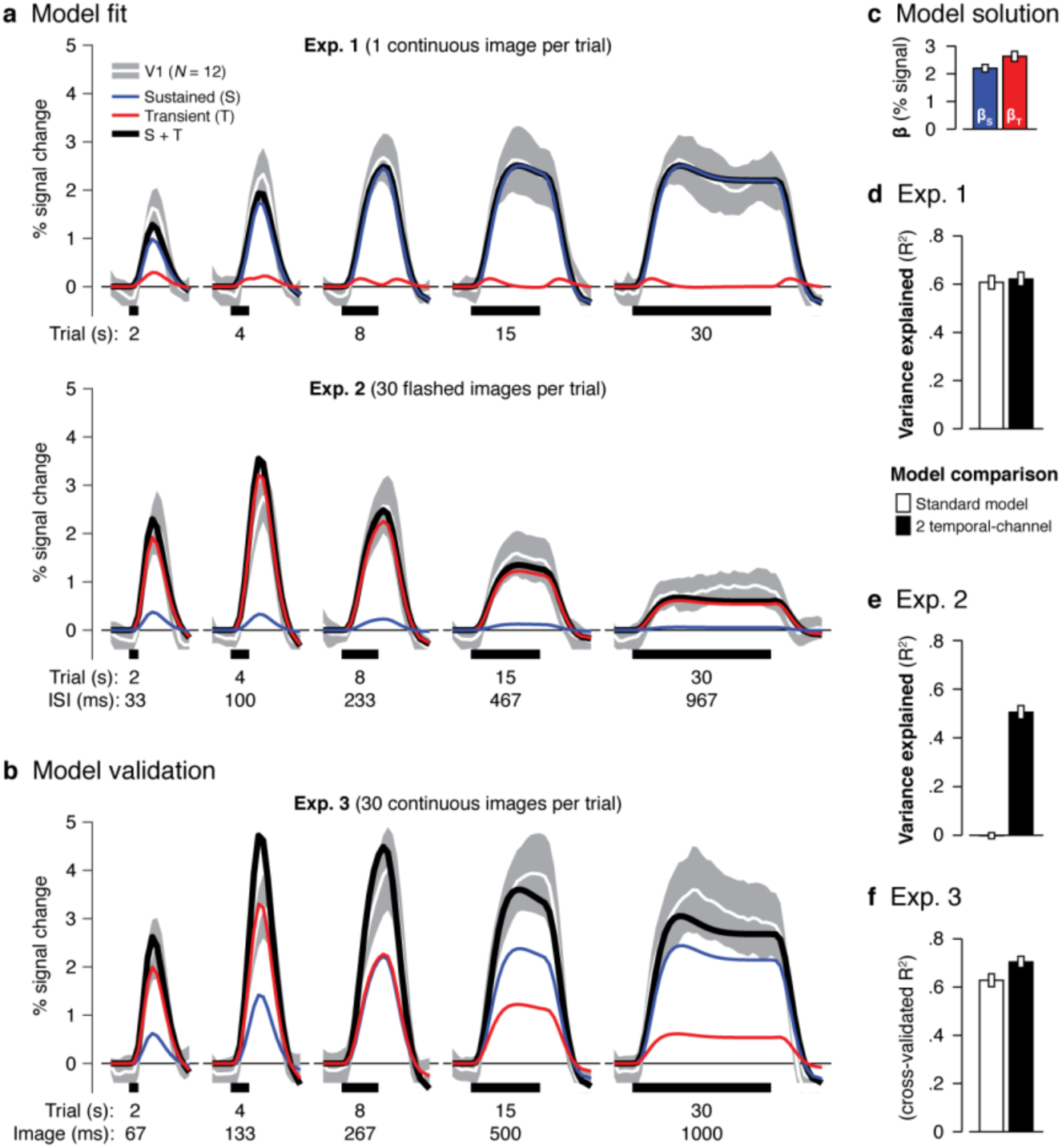
Sustained and transient contributions to V1 fMRI responses. (**a**) Measured V1 responses in Experiments 1 and 2 are plotted as the mean (*white*) ± 1 standard deviation (*gray*) across 12 participants for each trial duration. Superimposed are the predictions of the 2 temporal-channel model fit across data from both experiments. *Blue:* sustained predictor weighted by *β_s_*; *Red*: transient predictor weighted by *β_T_; Black:* prediction of the 2 temporal-channel model, which is the addition of the two channels, (**b**) Measured V1 responses and cross-validated model prediction for Experiment 3. The sustained and transient predictors are respectively weighted with *β_s_* and *β_T_* fitted from Experiments 1 and 2 [see (c)]. In all panels, trial durations are illustrated below the x-axis, and curves extend 2 s before the onset and 12 s after the offset each trial, (**c**) The model solution (*β_s_* and *β_T_*) for V1 fit with the two temporal-channel model using data concatenated across Experiments 1 and 2. **(d-f)** Comparison of the variance explained (*R^2^*) by 2 temporal-channel model vs. the standard model for each experiment. Error bars in (c–f) indicate ± 1 SEM across participants.

Second, the 2 temporal-channel model explains V1 responses to both Experiments 1 and 2, but the standard model fails to explain responses to transient stimuli in Experiment 2. That is, the 2 temporal-channel model fit to both experiments, explained 62% ± 3% (mean ± 1 standard error of the mean across participants, SEM) of V1 response variance in Experiment 1 (**Fig. 4d**) and 51% ± 3% of the variance in Experiment 2 (**Fig. 4e**). In contrast, the standard model fit to both experiments explained 61% ± 3% of the variance in Experiment 1 (**Fig. 4d**) but less than 1% ± 1% of the variance in Experiment 2 (**Fig. 4e**). Thus, while the standard model captured V1 responses to the long stimulus presentations in the first experiment, it failed to capture responses to transient stimuli in the second experiment (**Fig. 2a**).

Third, the 2 temporal-channel model with *β_S_* and *β_T_* fit from Experiments 1 and 2 (**Fig. 4c**) accurately predicted independent data from Experiment 3 (**Fig. 4b**). The sustained contribution (**Fig. 4a-b**, *blue*) in Experiment 3 was comparable to Experiment 1 (these experiments have the same total duration of stimulation per trial), and the transient contribution (**Fig. 4a-b**, *red*) in Experiment 3 was similar to Experiment 2 (these experiments have the same number of transients per trial). Since both temporal channels provided a significant contribution to V1 and the contributions of the two channels are additive, Experiment 3 responses were higher than both Experiments 1 and 2 across all trial durations. Analysis of cross-validated *R^2^* showed that the 2 temporal-channel model explained 71% ± 2% of variance in Experiment 3 (**Fig. 4f**) even though the channel weights were estimated from responses to independent data with different temporal characteristics. The cross-validated *R^2^* of the 2 temporal-channel model was also significantly higher than the standard model (*t*_11_ = 5.92, *p* < 0.001, paired *t*-test), which only explained 63% ± 3% of response variance in Experiment 3 (**Fig. 4f**). In general, the two-temporal channel model outperformed the standard model in all visual areas tested, with a larger improvement in subsequent areas compared to V1 (**Supplementary Figs. S1, S2).** Thus, the 2 temporal-channel model predicts fMRI responses to visual stimuli across a three-fold range of presentation durations ranging from tens of milliseconds to tens of seconds.

### Do Temporal Processing Characteristics Differ Across Intermediate Visual Areas?

We next examined hV4 and hMT+ responses to the time-varying visual stimuli in Experiments 1–3, as the competing theories make different predictions regarding the contributions of sustained and transient channels to these regions. hV4 and hMT+ illustrated distinct patterns of responses. Like V1, hV4 showed higher responses in Experiment 3 (30 continuous images per trial) than either Experiment 1 (1 continuous image per trial) or Experiment 2 (30 flashed images per trial). Different from V1, hV4 exhibited equal or stronger responses to the brief transient visual stimuli in Experiment 2 than the sustained single images in Experiment 1 (**Fig. 5a**). Different than both V1 and hV4, hMT+ exhibited close to zero evoked responses for the sustained stimuli in Experiment 1 (except for onset and offset responses that are visible in trials of 8 s and longer, **Fig. 5b**). However, hMT+ showed substantial responses for transient stimuli in Experiment 2 that were comparable to Experiment 3, which had both transient and sustained stimulation. Together, these data suggest differences in temporal processing across hV4 and hMT+.

**Figure 5:**
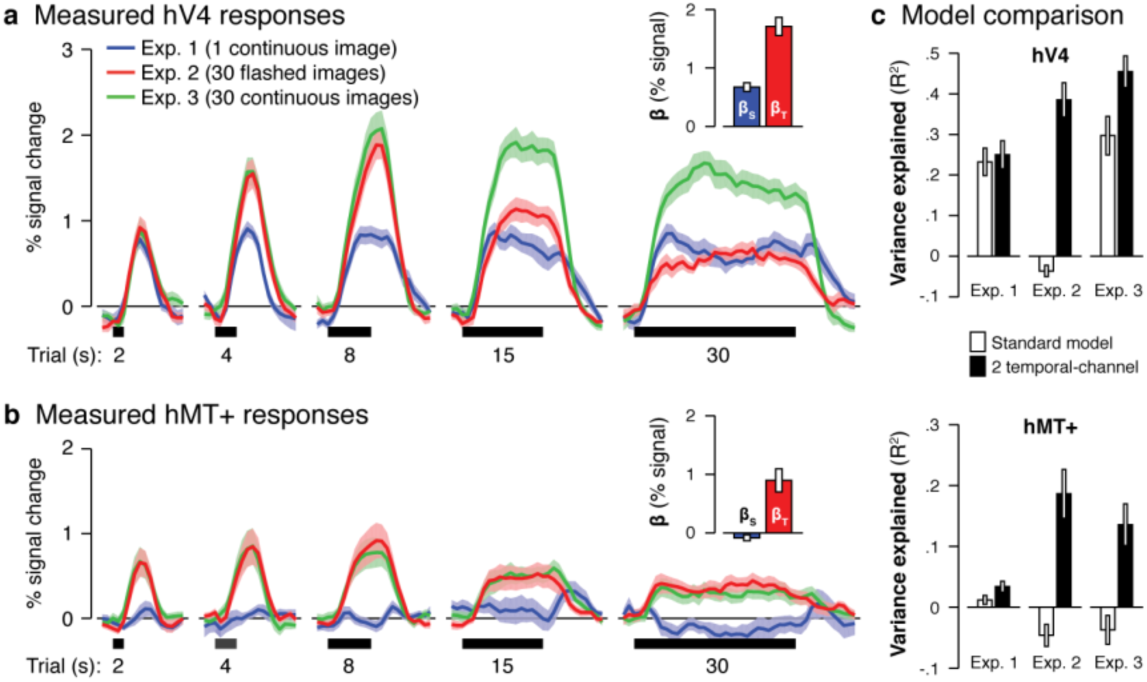
Differential sustained and transient contributions across hV4 and hMT+. (**a**) hV4 and (**b**) hMT+ responses for Experiment 1 (*blue*), Experiment 2 (*red*), and Experiment 3 (*green*). Curves show mean (solid line) ± 1 SEM across 12 participants (shaded area). Trial durations are indicated by the thick black lines below the x-axis, and curves extend 2 s before the onset and 12 s after the offset each trial. The model solution (*β_S_* and *β_T_*) for each region is plotted to as an inset with error bars representing ± 1 SEM across participants. (**c**) Comparison of the variance explained (*R^2^*) by 2 temporal-channel model vs. the standard model, both fit across data from Experiments 1 and 2. Model performance is quantified separately for each experiment both for hV4 (*top*) and hMT+ (*bottom*). Error bars indicate ± 1 SEM across participants.

Next, we quantified hV4 and hMT_+_ responses with the 2 temporal-channel encoding model. The model fits revealed that (i) in hV4 both channels contributed to responses, with the contribution of the transient channel about double that of the sustained channel (**Fig. 5a**, *inset*) and (ii) in hMT_+_ the transient channel substantially contributed to responses, but the sustained channel had close to zero contribution (**Fig. 5b**, *inset*). Across both regions, the 2 temporal-channel model fit data from Experiment 2 better than the standard model (*ts* > 4.06, *ps* < 0.01, paired *t*-test on *R^2^* values for each ROI) and also better predicted data from Experiment 3 than the standard model (*ts* > 4.18, *ps* < 0.05, paired *t*-test on cross-validated *R^2^* values for each ROI; **Fig. 5c**). These results demonstrate that not only does the 2 temporal-channel model perform significantly better than the standard model at intermediate stages of the visual hierarchy, but that the contributions of transient and sustained channels differ across hV4 and hMT_+_.

### What Is the Topology of Sustained and Transient Channels Across Visual Cortex?

To complement the ROI approach, we next visualized the spatial topology of the sustained and transient channels across visual cortex, which enables mapping channel contributions at the voxel-level.

Examining the contribution of sustained and transient channels across ventral and lateral occipito-temporal cortex revealed two main findings. First, lateral occipito-temporal cortex was devoid of contributions from the sustained channel, but had substantial contributions from the transient channel (**Fig. 6a**). This effect was widespread and included not only voxels in hMT_+_, as predicted by the prior analysis, but also extended (i) posteriorly into portions of lateral occipital areas LO-1 and LO-2, and (ii) ventrally into the inferior occipital gyrus and lateral fusiform gyrus. Dorsal regions along the intraparietal sulcus also showed negligible sustained responses (**Supplementary Fig. S3b**). Second, in ventral occipito-temporal cortex, regions along the posterior collateral sulcus and medial fusiform gyrus (where hV4, VO-1, and VO-2 are located) showed both transient and sustained responses, with larger contributions from the transient than sustained channel (**Fig. 6a**).

**Figure 6:**
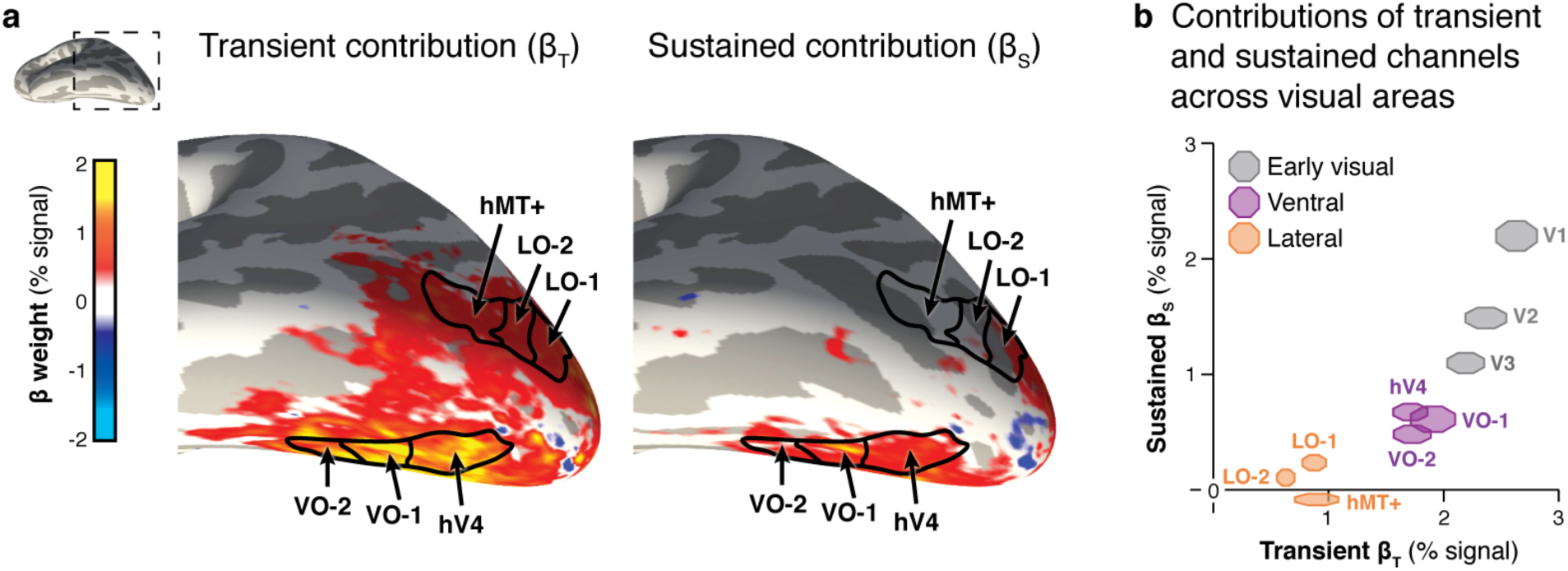
Differential transient and sustained contributions across ventral and lateral regions. (**a**) Ventrolateral view of occipito-temporal cortex (see inset) depicting group-averaged (*N* = 12) maps of the contributions of transient (*left*) and sustained (*right*) channels. We first estimated *β* weights of each channel in each voxel in each participant’s native brain space. *β* weight maps were transformed to the FreeSurfer average brain using cortex-based alignment and averaged across participants in this common cortical space. The resulting group maps were thresholded to exclude voxels with weak contributions (–0.1 > *β* > 0.1). Boundaries of ventral and lateral regions (*black*) are derived from the Wang Atlas, with hMT_+_ as the union of TO-1 and TO-2. (**b**) Contributions (*β* weights) of transient (x-axis) and sustained (y-axis) channels to each visual area as estimated by the 2 temporal-channel model. Marker size spans ± 1 SEM across 12 participants in each dimension and *β* weights were solved by fitting the 2 temporal-channel using data concatenated across Experiments 1 and 2.

We quantified the mean contributions of transient and sustained channels across visual areas spanning occipito-temporal cortex. Our results showed differences in the contributions of sustained and transient channels across early visual cortex (V1–V3), ventral occipito-temporal cortex (hV4, VO-1, and VO-2), and lateral occipito-temporal cortex (LO-1, LO-2, and hMT+; **Fig. 6b**, significant temporal-channel by cluster interaction, *F*_2,_ 22 = 13.16, *p* < 0.001, two-way ANOVA on *β* weights with factors of temporal-channel [sustained/transient] and visual cluster [early/ventral/lateral]). From V1 to higher-order areas, there was a larger drop in the contribution of the sustained channel than the transient channel (**Fig. 6b**). Nevertheless, there were significant differences among clusters: in ventral areas both sustained and transient channels contributed to responses, but in lateral areas responses were dominated by the transient channel.

The spatial topology of sustained and transient channels also revealed differences in temporal processing within regions. Specifically, in early visual cortex (V1-V3) the sustained channel was robust in eccentricities < 20°, but declined in more peripheral eccentricities (**Fig. 7a**, *right*). In contrast, the transient channel contributed to responses across a larger range of eccentricities that extend further into the periphery (> 20°; **Fig. 7a**, *left*). We quantified these effects by measuring the contributions of the 2 temporal-channels across eccentricities using uniformly-sized disc ROIs defined along the horizontal meridian representations in V1 and V2/V3 (**Fig. 7b**). This quantification showed that in early visual areas the magnitude of the sustained channel declined more rapidly with eccentricity than the transient channel, to the extent that at eccentricities of 40° there still was a 0.90 ± 0.17% transient response but less than 0.26 ± 0.07% of a sustained response (**Fig. 7b**). Further, the decline of the sustained channel with eccentricity occurred more rapidly in V2/V3 than V1. Together, we find a differential contribution of transient and sustained channels across eccentricities and areas (significant three-way interaction of temporal channel [sustained or transient], visual area [V1 or V2/V3], and eccentricity [5°, 10°, 20°, or 40°], *F*_3, 33_ = 3.18, *p* < 0.05, three-way ANOVA on the *β* weights).

**Figure 7:**
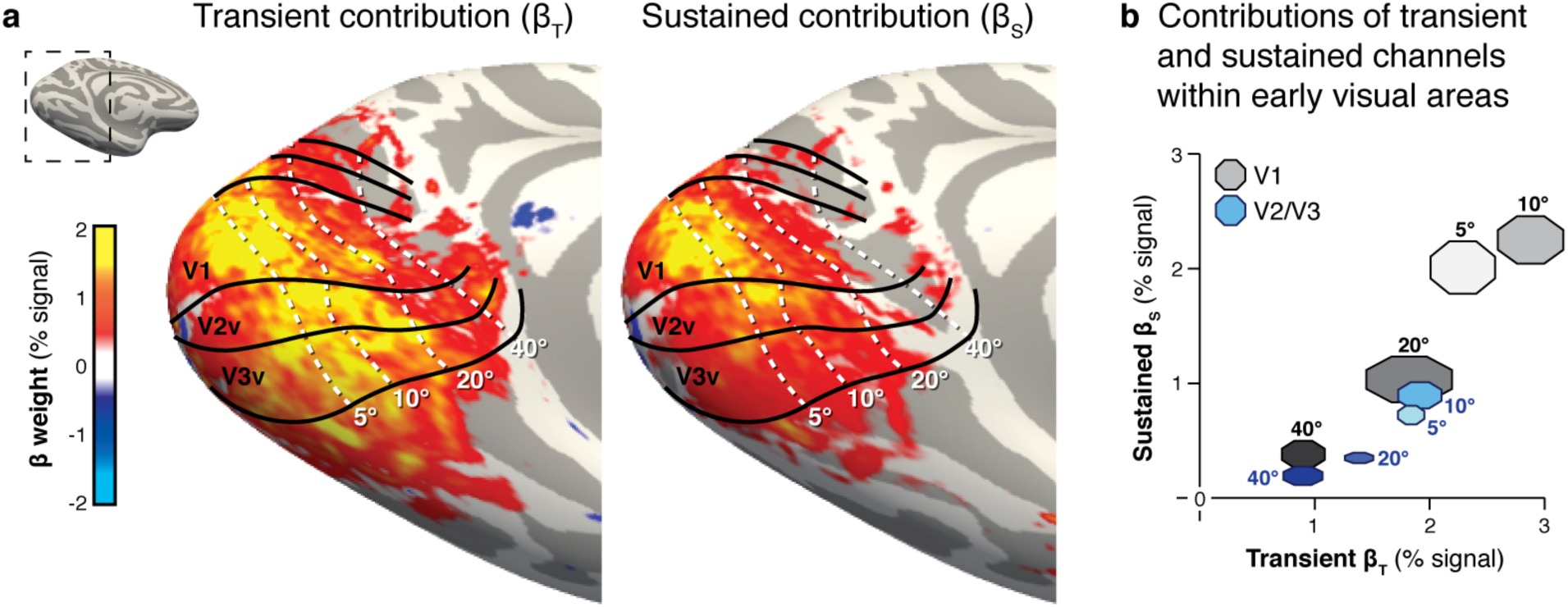
Differential transient and sustained contributions across central and peripheral eccentricities. (**a**) Medial cortical surface zoomed on the occipital lobe (see inset) depicting group-averaged (*N* = 12) maps of the contributions of transient (*left*) and sustained (*right*) channels. We first estimated *β* weights of each channel in each voxel in each participant’s native brain space. *β* weight maps were transformed to the FreeSurfer average brain using cortex-based alignment and averaged across participants in this common cortical space. The resulting group maps were thresholded to exclude voxels with weak contributions (–0.1 > *β* > 0.1). Regional boundaries (*black*) and eccentricity bands (*white*) of early visual areas are derived from the Benson Atlas. (**b**) Contributions of transient (x-axis) and sustained (y-axis) channels across eccentricities along the horizontal representation in V1 (grays) and V2/V3 (blues) as estimated by the 2 temporal-channel model. Eccentricities range from 5° (*lightest markers*) to 40° (*darkest markers*). Marker size spans ± 1 SEM across 12 participants in each dimension and *β* weights were solved by fitting the 2 temporal-channel using data concatenated across Experiments 1 and 2.

## DISCUSSION

Our results show that a 2 temporal-channel model of neural responses, containing sustained and transient channels, is a parsimonious encoding model that predicts fMRI responses in human visual cortex to visual stimuli across a broad range of durations, from tens of milliseconds to tens of seconds. Critically, our data address the ongoing debate regarding the contribution of sustained and transient channels in extrastriate cortex. Consistent with the prevailing view^2, 6 7^, we find that the transient channel dominates hMT+ responses and peripheral eccentricity representations of V1^36^. Importantly, we show for the first time that this temporal processing characteristic extends to human lateral occipito-temporal cortex as well as peripheral eccentricity representations of V2 and V3. In contrast to the prevailing view, we find that both sustained and transient channels drive responses not only in hV4 but also ventral occipito-temporal regions VO-1 and VO-2, with a surprisingly larger contribution of the transient than sustained channel. This finding argues against the view that the ventral stream primarily codes static visual information and suggests a rethinking of the role of transient processing in the visual system.

### Differential Transient and Sustained Responses Across Visual Cortex

Our research fills a large gap in knowledge regarding temporal processing in human visual cortex by showing that (i) the 2 temporal-channel model is applicable to at least 10 additional visuals areas beyond V1^30-32, 36, 38^, and (ii) temporal processing is a key functional attribute that differentiates visual areas.

Our observation that hMT_+_ responses to sustained stimuli are close to zero is consistent with the prevailing view that (i) hMT_+_ is involved in processing visual dynamics rather than static information, and that (ii) inputs to hMT_+_ are M dominated^7^. Notably, we found that neighboring regions, LO-1 and LO-2, also have close to zero sustained responses. This finding is interesting because LO-2, which is thought to be involved in visual processing of objects^39, 40^ and bodyparts^41^, shows more robust responses to rapidly presented stimuli compared to nearby category-selective regions^42^. The present data suggest that this characteristic may be an outcome of a dominant transient channel – a hypothesis that can be tested in future research.

Inconsistent with the prevailing view, we found that hV4 showed not only sustained responses as expected^8, 10, 24, 43^, but also large transient responses. While this observation is consistent with reports that macaque V4 receives both P and M inputs^5, 9^, it is unexpected that the transient contributions to hV4 exceed the sustained contributions. Further, unlike the macaque brain, where portions of V4 are adjacent to MT, hV4 is ~3 cm away from hMT_+_, indicating that this finding cannot be explained by the proximity of these two regions. While the sustained channel is often associated with coding static visual input and the transient channel with coding visual dynamics^10, 11, 44^, visual transients can also indicate changes to the content of the visual input. Indeed, in our experiments, transients occurred when stimuli changed (i.e., when a new image was shown or an image was replaced by a uniform gray screen). Since the function of the ventral stream is to derive the content of the visual input, the fast transient visual processing we observe in ventral stream regions may enable rapid processing of visual changes^45^, which in turn, may foster detection of novel stimuli and rapid extraction of the gist of the visual scene.

It is interesting that temporal processing in lateral occipito-temporal regions, like that of far peripheral eccentricities (> 20°) in early visual cortex, was dominated by the transient channel, and temporal processing in ventral occipito-temporal regions, like lower eccentricities in early visual cortex, showed a dual channel contribution. These functional characteristics may be anatomically supported by white matter connections from peripheral representations of early visual areas to MT and nearby regions^46^ that are separate from white matter connections from central representations of early visual cortex to ventral occipito-temporal regions^47^. Furthermore, the diminished sustained responses in the periphery of early visual cortex is consistent with prior findings showing reduced P inputs^48^ and diminished sustained luminance responses^36^ in peripheral compared to central V1, as well as faster perception in the periphery^49^.

### Implications for Modeling fMRI Signals: Millisecond Timing Matters

Our data have important implications regarding modeling fMRI signals and understanding temporal processing in the human brain because they show that (i) varying the temporal characteristics of the visual stimulus in the millisecond range has observable effects on fMRI responses in the second range, and (ii) by considering the contribution of a transient neural channel, the encoding model can account for nonlinearities in fMRI responses for rapid and short visual stimuli^19-21,23, 50^.

Our data extend the original linear model of fMRI signals^19, 25^ by showing the importance of modeling the temporal properties of neural responses at millisecond resolution to accurately predict fMRI signals. In their original study, Boynton et al.^19^ noted consistent deviations from the linear (standard) model in short durations (3–6 s) in which the model underestimated fMRI signals. These nonlinearities are exacerbated in experiments using even shorter stimuli (1/4 to 2 s^21, 23, 50^). Boynton and colleagues suggested that neural adaptation or transient responses may explain deviations from linearity. We favor the interpretation that transient responses account for nonlinearities for two reasons: (i) Taking into account the neural transient channel resolves this nonlinearity and can predict not only our measurements but also prior data showing nonlinearities^23^ (**Supplementary Fig. S4**). (ii) Adaptation would have resulted in declining responses during long trials of continuous presentation of a single stimulus^45^. However, we observed negligible adaptation in striate (e.g. V1) and extrastriate (e.g. hV4) areas even during the 30 s single continuous image trials.

It is worthwhile noting that while the 2 temporal-channel model provides a significant improvement in modeling fMRI signals, our model does not explain the entire variance of fMRI signals (**Supplementary Fig. S2**) and it does not account for all nonlinearities, as it still overestimates the responses in extrastriate regions during short trials in Experiment 3 (**Supplementary Fig. S1**). A recent encoding approach^51^ addressed these types of nonlinearities by implementing compressive temporal summation (CTS) that predicts sub-additive responses, together with dynamic normalization that depends on response history. Therefore, an important direction for future research would be to add CTS and dynamic normalization to each of the temporal channels, to test if such a combined approach further improves predictions of temporal dynamics in visual cortex, especially in higher-order areas, which have larger compressive temporal summation^51^ and larger adaption^43, 52^. Another direction for future research could combine the 2 temporal-channel model with a spatial receptive field model^26-29^ to generate a complete spatio-temporal understanding of visual responses.

Given the pervasive use of the standard linear model in fMRI research, our results have broad implications for fMRI studies in any part of the brain. We find that timing of stimuli in the millisecond range has a large impact on the magnitude of fMRI responses, which has important implications for interpreting results of studies that vary the temporal characteristics of stimuli across conditions (e.g.^53^). Critically, we demonstrate that rather than ignoring fast cortical processing because of nonlinearities in the standard model, it is possible to generate neural predictions at sub-second resolution and use them to accurately predict fMRI responses (see also^51^). These new encoding approaches open exciting new opportunities for investigating fast cortical mechanisms using fMRI in many domains including somatosensory, auditory, and high-level cognitive processing.

In sum, our experiments elucidate the characteristics of temporal processing across human visual cortex. These findings are important because they (i) explicate for the first time the contribution of transient and sustained visual responses across human visual cortex beyond V1, and (ii) show that accounting for neural responses in the millisecond range has important consequences for understanding fMRI signals in the second range in any part of the brain.

## ACKNOWLEDGEMENTS

This research was supported by NEI grant 1R01EY02391501A1. We thank Justin Gardner, Kendrick Kay, Anthony Norcia, Brian Wandell, and Kevin Weiner for fruitful discussions and constructive feedback.

## ONLINE METHODS

### Participants

Twelve participants (6 males, 6 females) with normal or corrected-to-normal vision participated in this study. All participants provided written informed consent, and the experimental protocol was approved by the Stanford University Institutional Review Board. Each individual participated in three fMRI sessions, two used to fit and validate the 2 temporal-channel model, and one session in which we conducted population receptive field (pRF) mapping^26^ to define retinotopic cortical regions and another experiment to define human motion-sensitive area (hMT_+_)^54-56^.

### Temporal Channels Experiments

#### Visual stimuli

We used full-field phase-scrambled stimuli to generate robust visual responses while minimizing cognitive factors. Stimuli consisted of grayscale images that were generated by randomizing the phase spectrum of naturalistic images used in our previous publications^42^ (**Fig. 1a**, *right*). We normalized the mean luminance and distribution of grayscale values in each image using the SHINE toolbox^57^. Stimuli were displayed to participants in the scanner using an Eiki LC-WUL100L projector (resolution: 1920 × 1200; refresh rate: 60 Hz) that was controlled by an Apple MacBook Pro using MATLAB (http://www.mathworks.com) and functions from Psychophysics Toolbox^58^ (http://psychtoolbox.org). Participants viewed the projected images through an auxiliary mirror mounted on the RF coil. The mirror was adjusted in each participant such that stimuli spanned approximately 40° of visual angle in the vertical dimension and 60° of visual angle in the horizontal dimension.

#### Experimental design

To obtain data that can be used to estimate and test the 2 temporal-channel encoding model, we introduce a novel fMRI paradigm that estimates independent sustained and transient contributions to fMRI responses across visual cortex using three experiments. All three experiments used the same stimuli, trial durations, and task, and only varied in their temporal presentation of the stimuli as detailed below and illustrated in Fig. 1.

##### Experiment 1 largely sustained stimulation

phase-scrambled images were shown in trials of varying durations (2, 4, 8, 15, or 30 s per trial) in which a single phase-scrambled image was shown for the entire duration of the trial (Fig. 1, *blue*). Before and after each trial there was a 12 s baseline period (blank gray screen matched to the mean luminance of the stimuli). Across trials the number of stimuli (one per trial) and transients (at the onset and offset of each stimulus) are matched; just the duration of sustained stimulation varies. This experiment was designed to primarily activate the sustained channel, especially in the long trials.

##### Experiment 2 argely transient stimulation

used the same trial durations and general experimental design as Experiment 1, except that in each trial 30 different phase-scrambled images were shown briefly, each for 33 ms. Thus, the number of stimuli, number of transients, and total duration of visual stimulation are matched across trial durations. The only factor that varied across trials was the inter-stimulus interval (ISI) between consecutively presented images. The ISI consisted of a blank mean-luminance screen that was 33 ms long in the 2 s trials, 100 ms in the 4 s trials, 233 ms in the 8 s trials, 467 ms in the 15 s trials, and 967 ms in the 30 s trials (Fig. 1, *red*). This experiment was designed to maximally drive the transient channel and minimally the sustained channel since each image was shown for only 33 ms.

##### Experiment 3 combined sustained and transient stimulation

used the same design as Experiment 2, except that in each trial we presented 30 different phase-scrambled images in a continuous fashion without an ISI between sequential images. The durations of images (67, 133, 267, 500, or 1000 ms per image) varied across trials that were matched in length to Experiments 1, whereby the 67 ms presentations occurred in the 2 s trials and the 1000 ms presentations occurred in the 30 s trials (**Fig. 1**, *green*). This experiment was designed to drive both the sustained and transient channels: (i) during the entire trial duration there was always a stimulus on the screen and (ii) there were always 30 different images per trial.

##### Task

In all three experiments, participants were instructed to fixate on a small, central dot, and respond by button press when it changed color (occurring randomly once every 2–14 s, 8 s on average).

Experiments were designed such that if the sustained component is dominant, then Experiments 1 and 3 should yield similar responses since they have the same overall duration of stimulation. However, if the transient component is dominant then responses in Experiments 2 and 3 should be similar as they have the same number of transients. Finally, if transient and sustained channels contribute independently to responses, then fMRI signals in Experiment 3 that has both types of visual stimulation should be higher than either Experiments 1 or 2.

#### Data acquisition

Data were obtained with a TR of 1 s and a surface coil, collecting 16 slices per acquisition. To gain full coverage of occipito-temporal cortex, all participants completed two scan sessions on different days with partially overlapping slice prescriptions. In each session participants viewed three 288 s runs of each experiment. Each run of each experiment contained of two repeats of each trial duration presented in random order. Data from different sessions were pooled in each participant’s volume anatomy. The three runs of each experiment were blocked in each participant, and the order of experiments was counterbalanced between participants.

*Population receptive field (pRF) mapping:* To delineate retinotopic boundaries, we collected four 200 s runs of pRF mapping in each participant, same as in Dumoulin and Wandell^26^. In this experiment, a bar swept across a circular aperture (40° by 40° of visual angle) in eight directions and baseline periods were interspersed throughout each run. Participants performed the same color exchange fixation task as in the main experiments. We used the data to generate polar angle and eccentricity maps, which were used to define retinotopic visual areas as in our prior publications^41,42, 59^.

*hMT_+_ localizer:* To functionally define hMT_+_, in each participant we collected one 300 s run of a motion localizer experiment^41^. In this experiment, low contrast, 40° by 40°, concentric rings alternated between 16 s periods of motion (expansion/contraction) and 16 s periods of a stationary display. The experiment contained 6 cycles of alternating moving and stationary trials. Participants performed the color exchange fixation task.

#### Magnetic Resonance Imaging (MRI)

MRI data were collected using a 3T GE Signa MR750 scanner at the Center for Cognitive and Neurobiological Imaging (CNI) at Stanford University.

##### fMRI

We used a Nova 16-channel visual array coil (http://novamedical.com) to give participants a large unobstructed visual field of view. In each participant, we acquired two partially-overlapping oblique slice prescriptions in separate scan sessions that together fully cover occipito-temporal cortex (resolution: 2.4 × 2.4 × 2.4 mm; one-shot T2*-sensitive gradient echo acquisition sequence: FOV = 192 mm, TE = 30 ms, TR = 1000 ms, and flip angle = 73°). We also collected T1-weighted inplane images with the same prescription as the functional data to align each participant’s data to their high-resolution whole brain anatomy.

In a separate session, we obtained pRF mapping and hMT_+_ localizer data with the same RF coil setup and spatial resolution using 28 oblique slices covering the same brain volume but with a longer TR (resolution: 2.4 × 2.4 × 2.4 mm; one-shot T2^*^-sensitive gradient echo acquisition sequence: FOV = 192 mm, TE = 30 ms, TR = 2000 ms, and flip angle = 77°). We again collected T1-weighted inplane images in the same prescription to finely align inplane data to the whole brain anatomy of each participant.

##### Anatomical MRI

We acquired a whole-brain, anatomical volume in each participant using a Nova 32-channel head coil (resolution: 1 × 1 × 1 mm; T1-weighted BRAVO pulse sequence: TI = 450 ms, flip angle = 12°, 1 NEX, FOV = 240 mm).

**Data analysis**

Data were analyzed with MATLAB using code from vistasoft (http://github.com/vistalab) and FreeSurfer (http://freesurfer.net).

#### Data pre-processing

Functional data were aligned to each participant’s native anatomical space using T1-weighted inplane images, and volumes acquired within the first 8 s of each run were discarded to allow time for magnetization to stabilize. We then performed slice time correction, motion compensation (within and between scans), and transformed voxel time series to units of percent signal change. To normalize the baseline level of response across experiments, we subtracted from time points in each run the mean signal across the 4 s periods preceding the trial onsets in each run. This baseline removal procedure centers the mean response for the blank screen around zero to improve cross-validation performance^60^ and to enable comparison of trial responses relative to the blank baseline.

#### Region of interest (ROI) definition

Areas V1, V2, V3, V3A, V3B, hV4, VO-1, VO-2, LO-1, and LO-2 were defined in each participant’s native anatomical space using data from the pRF mapping experiment as in prior publications^41, 42, 59^ (**Supplementary Fig. S2a**). To improve model performance in later visual areas, we fit one pRF to the run-averaged time series of each voxel using the compressive spatial summation (CSS) variant of the standard pRF model^61, 62^. ROIs were drawn bilaterally on each participant’s cortical surface using the resulting polar angle and eccentricity maps. We excluded from ROI analyses voxels with pRF fits that explain less than 5% of their response variance. The distance between hV4 and hMT_+_ was calculated in each hemisphere by finding the centroid of each region and computing the Euclidean distance between the two points on the brain volume (mean distance = 3.04 cm, standard deviation = 1.24 cm). Dorsal visual areas V3A and V3B were also defined in each participant, but here we focus on regions in early visual cortex (V1, V2, and V3), ventral occipito-temporal cortex (hV4, VO-1, and VO-2), and lateral occipito-temporal cortex (LO-1, LO-2, and hMT_+_) because these regions have been more widely studied with regard to their temporal capacity than dorsal stream regions^7, 9 24, 30-32, 63^. Data for V3A and V3B are included in **Supplementary Fig. S3**.

We defined hMT_+_ bilaterally in each participant using data from the motion localizer experiment as described in our previous publications^41^. hMT_+_ was defined as voxels in the posterior inferior temporal sulcus^64^ that responded significantly (*t* > 3) more to moving than stationary stimuli.

#### 2 temporal-channel model

In typical analysis of fMRI responses^19, 25^, the stimulus vector is convolved with the hemodynamic response function (HRF) to obtain a prediction of the fMRI response. However, this model does not account for distinct temporal channels of neural responses^30-32, 65^. To generate predicted fMRI responses accounting for the temporal channels, we implemented an encoding approach similar to Horiguchi^36^. The code used to implement the encoding model is available for download (https://github.com/VPNL/TemporalChannels).

The model illustrated in **Fig. 3** shows the procedure. First, we estimate the neural response of each channel by convolving the stimulus (**Fig. 3a**) separately with the neural impulse response function (IRF) for the sustained channel (**Fig. 3b,** *blue channel IRF*) and the transient channel (**Fig. 3b,** *red channel IRF*). This generates the predicted neural response to the visual stimulus for each channel. Then, the estimated neural responses for each channel are convolved with the hemodynamic response function (HRF, **Fig. 3c**) and summed to generate a prediction of the fMRI response. We use a general linear model (GLM) to solve for the contributions of the sustained and transient channels (*β* weights) given the measured fMRI responses.

The *sustained neural channel* is characterized by a monophasic IRF_S_ that generates a response for the entire duration of a stimulus. The *transient neural channel* is characterized by a biphasic IRF_T_ that generates a brief response at the onset and offset of an image^30-32, 34, 35^. The transient channel also contains a nonlinearity (squaring operation), that generates positive responses both from the onset and offset of the stimulus, as firing rates associated with transient ‘on’ or ‘off’ responses are positive^66^ and metabolically demanding^36, 37^. The nonlinearities in this model are at the neural level, and a linear relationship is assumed between the neural and BOLD responses.

*Modeling the neural impulse response:* Our model used impulse response functions estimated from human psychophysics^35^ (**Fig. 3b**) to approximate the temporal sensitivity of the human visual system. These IRFs are expressed as the difference between excitatory and inhibitory linear filters. The excitatory filter is expressed as

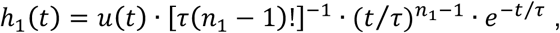

where *u*(*t*) is the unit step function at time *t; τ* is a fitted time constant, and *n*_1_ is the number of stages in the excitatory filter. The inhibitory filter incorporates the same time constant and is expressed as

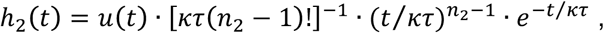

where *κ* is the ratio of time constants for the two filters and *n*_2_ is the number of stages in the inhibitory filter. Both the sustained and transient channel IRFs are derived with the formula

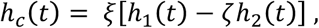

where the normalization parameter *ξ* is used to match the height of the functions and is equal to 1 for IRF_S_ and 1.44 for IRF_T_; the transience parameter *ζ* is equal to 0 for IRF_S_ and 1 for IRF_T_.

The other parameters are taken directly from Watson^35^ and are: *τ* = 4.94 ms, *κ* = 1.33, *n*_1_ = 9, and *n_2_* = 10.

##### Modeling the visual input

Since the neural impulse response to a stimulus occurs on a millisecond timescale, we code each stimulus sequence in milliseconds. The stimulus is coded as a binary vector of ones and zeros, where one represents the presence of a stimulus and zero indicates when there is no stimulus, just a blank mean luminance screen (**Fig. 3a**). To capture the digital transitions of the display (constrained by the 60 Hz refresh rate of the projector), a 17 ms gap is coded at the offset of each image. Next, the stimulus vector is convolved separately with each channel IRF to generate separate sustained and transient neural response predictors (**Fig. 3b**). To model the corresponding fMRI responses from each channel, each of the two neural response predictors are convolved with a HRF (**Fig. 3c**) that was sampled at the same high (millisecond) temporal resolution of the neural response predictors. Here, we slightly adapted the parameters of the canonical HRF implemented in SPM8 (http://www.fil.ion.ucl.ac.uk/spm/software/spm8) to better capture the rise and fall of the BOLD response in our measurements (delay of peak response = 5 s, delay of undershoot = 14 s, kernel length = 28 s).

##### Fitting the 2 temporal-channel model

Since the HRF acts as a low-pass temporal filter, this enables us to resample the predicted fMRI response to the lower temporal resolution of the acquired fMRI data (TR = 1 s). This resampled fMRI response predictor is compared to measured fMRI responses to solve for the contributions (*β* weights) of each channel. We normalized the predicted fMRI responses across the two channels, such that the maximal height is the same across both channels. Then we used a GLM to estimate the weights of the sustained (*β_S_*) and transient (*β_T_*) predictors by comparing the predicted responses to the measured response using data across all runs of Experiments 1 and 2. The GLM is applied to the mean response of each visual area in each participant. Quantification of model performance in each of Experiments 1 and 2 is presented in **Fig. 4d-e** for V1, in **Fig. 5c** for hV4 and hMT_+_, and in **Supplementary Fig. S2b-c** for all ROIs. The predicted fMRI responses generated by the model are shown in **Fig. 4a** for V1 and **Supplementary Fig. S1** for other ROIs.

##### Validating the 2 temporal-channel encoding model

We assessed the predictive power of the 2 temporal-channel model by testing how well it predicts responses in independent data obtained in Experiment 3 (**Fig. 4b**). Thus, we coded the visual stimulation of Experiment 3 in the same manner described above, and convolved it separately with the IRFs of the sustained and transient neural channels to generate the neural predictors. These neural predictors were then convolved with HRF and downsampled to 1 s. Then we multiplied each channel’s fMRI response predictor with its respective *β* weight (*β_S_* or *β_T_*) that was estimated with data concatenated across Experiments 1 and 2. We then tested how well the predicted responses matched the measured response in Experiment 3, operationalized as cross-validated *R^2^,* also known as the coefficient of determination. That is, the proportion of response variance explained using *β* weights that were estimated from independent data. Although conceptually like the classical *R^2^* statistic, cross-validated *R^2^* can be negative when the residual variance of an inaccurate prediction exceeds the variance in the measured response. Quantification of cross-validation performance is shown in **Fig. 4f** for V1, in **Fig. 5c** for hV4 and hMT+, and in **Supplementary Fig. S2d** for all ROIs. The predicted fMRI responses generated by the model are shown in **Fig. 4b** for V1 and **Supplementary Fig. S1** for other ROIs.

##### Fitting and validating the standard model

For model comparison to the standard approach used in fMRI, we also fit a standard GLM to the data. The standard GLM predicts fMRI responses by convolving a stimulation vector with the HRF (basically steps **a** and **c** in **Fig. 3**). To describe the visual stimulation in our experiments, we used the same millisecond resolution visual stimulation vector as in the 2 temporal-channel model described above. After convolving with the same HRF used in the 2 temporal-channel model, we downsampled the predicted response to 1 s resolution to compare to measured fMRI responses. As the stimuli in all experiments were identical and the only difference was the timing, we generated a single predictor across Experiments 1 and 2 and fit one *β* weight. Like the 2 temporal-channel model, we fit the standard model to data concatenated across all runs of Experiments 1 and 2, and we then cross-validated this *β* weight on Experiment 3 data. Standard model predictions for V1 are shown in **Fig. 2a** and for other ROIs in **Supplementary Fig. S1**. Model performance is quantified separately for Experiments 1 and 2 in **Supplementary Fig. S2b-c**.

##### Model comparison

We used repeated measures analysis of variance (ANOVA) to compare the performance of the 2 temporal-channel model to the standard model across the visual hierarchy. To test for differences in the predictive power of the two models at various stages, we performed a two-way repeated measures ANOVA with factors of model (2 temporal-channel/standard) and visual field map cluster (early/ventral/lateral) on the cross-validated *R^2^* values from each ROI (see **Supplementary Fig. S2d** for ANOVAs including dorsal regions). For the 2 temporal-channel model, we additionally tested for differences in the relative contributions of the sustained and transient channels across the hierarchy using a two-way repeated measures ANOVA with factors of temporal channel (sustained/transient) and visual field map cluster (early/ventral/lateral) on the channel *β* weights estimated for each ROI using data from Experiments 1 and 2 (**Fig. 6b**).

##### Noise ceiling calculation

To compare the level of noise in measurements from different brain regions, we estimated the noise ceiling of each ROI using a procedure proposed by Kay^61^. This method estimates the maximal *R^2^* that a model could achieve given the level of noise in the data by simulating noisy measurements of a signal (with noise characteristics matched to the data) and using a bootstrapping procedure to estimate the median accuracy with which any given model could explain the simulated measurements. This procedure was performed separately for the 2 temporal-channel model and the standard model, and we plot the average noise ceiling estimate across both models for each ROI in **Supplementary Fig. S2**.

##### Generating group parameter maps of temporal channel contributions

To map the topology of contributions from the transient and sustained channels across visual cortex, in each participant we fit the 2 temporal-channel model in each voxel using data from Experiments 1 and 2, as described above. We then generated in each participant’s brain parameter maps of each temporal channel’s *β* weights. To generate group maps, we first registered all participants’ anatomy to the FreeSurfer average brain template using cortex-based alignment (CBA) in FreeSurfer version 5.3.c^67-70^. Then, we averaged these maps across participants on the common fsaverage cortical surface to obtain group maps (**Figs. 6a**, **7a**; **Supplementary Fig. S3b**). To independently validate our results from ROI analyses, group hV4, VO-1, VO-2, LO-1, LO-2, and hMT_+_ are outlined on the ventrolateral surface using the atlas developed by Wang and colleagues^71^ (**Fig. 6b**, **Supplementary Fig. S3b**). Note that hMT_+_ is defined based on the union of TO-1 and TO-2 maps^56^ from the Wang atlas^71^. To illustrate channel contributions in relation to the eccentricity map in early visual cortex, we used atlases developed by Benson and colleagues^72^ to overlay the regional boundaries of V1, V2, and V3 and trace eccentricity bands on the group average (**Fig. 7a**).

##### Measuring temporal channel contributions across eccentricities

To quantify changes in the contributions of transient and sustained channels within regions in early visual cortex as a function of eccentricity, we defined a series disc ROIs (each ~1 cm in diameter) in each hemisphere of the fsaverage cortical surface using the tksurfer package included with FreeSurfer. Using the Benson atlas^72^, uniformly-sized ROIs were dilated around mesh vertices centered on eccentricities of either 5°, 10°, 20°, or 40° (**see Fig. 7b**) both along the horizontal meridian representations in V1 and along the ventral and dorsal borders of V2/V3. Note that: (i) we used the Benson^72^ atlas as it contains eccentricities further into the periphery that were not mapped in pRF mapping experiment (visual angle was limited to 20° from fixation, see above), and (ii) we placed disk ROIs along the horizontal meridian as the phase-scrambled stimuli used in Experiments 1–3 extended further into the periphery in the horizontal than the vertical dimension (30° vs. 20° degrees from fixation, respectively). The disc ROIs were then transformed to each participant’s native brain space, and we fit the two temporal-channel model to data from Experiments 1 and 2 in each ROI as described previously (**Fig. 7b**). To compare differences in temporal channel contributions across eccentricities between V1 and V2/V3, we used a three-way, repeated measures ANOVA with factors of channel (transient or sustained), visual area (V1 or V2/V3), and eccentricity (5°, 10°, 20°, or 40°).

##### Explaining previously reported nonlinearities for short stimuli

To externally validate the 2 temporal-channel model, we simulated the temporal characteristics of visual stimulation in a previous study by Birn et al.^23^ that measured V1 responses to brief presentations of a checkerboard stimulus that was contrast inverted at a rate of 8 Hz (see **Supplementary Fig. S4a**). We used data from the Birn study because it showed other experimental conditions in which the standard model underestimated measured fMRI responses using stimulus presentations in the millisecond range. To predict responses to this 8 Hz stimulus with our model, we coded each 125 ms phase of a flicker sequence as a separate image within a trial (as described for Experiment 3). Using this coding of the stimulus in the Birn study, we used our 2-temporal channel model to generate fMRI response predictors of each channel, and multiplied the predicted response of each channel using the *β* weights estimated for V1 across Experiments 1 and 2 (**Supplementary Fig. S4b**, *left*). Finally, we used the same stimulus model and the *β* weight estimated by the standard model for V1 (**Supplementary Fig. S4b**, *right*) to compare the validity of the two models.

---

**Supplementary Figure S1:**
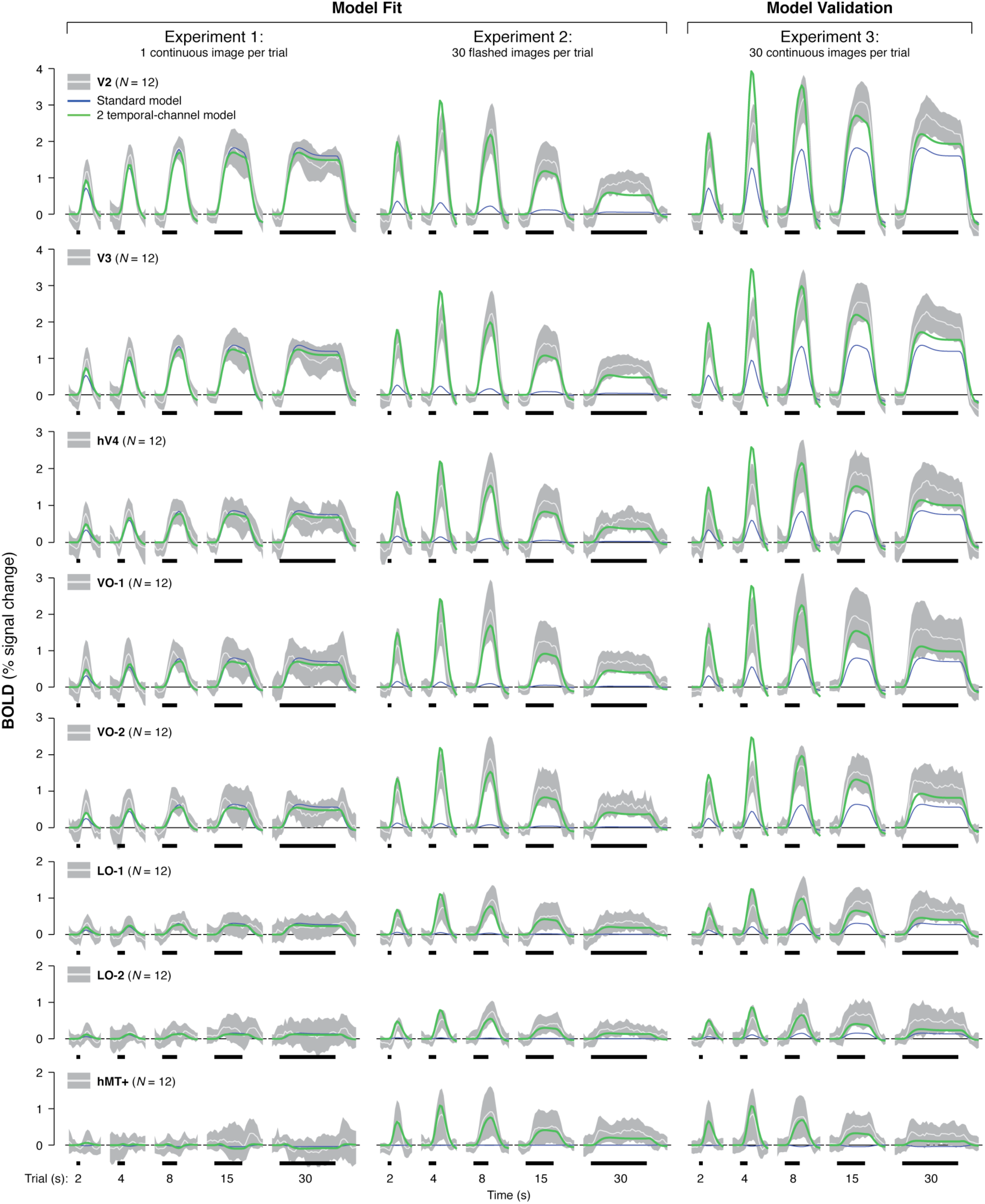
Modeling sustained and transient contributions across the visual hierarchy. Each row depicts the average ROI response (*white*) ± 1 standard deviation (*gray*) across 12 participants. *Left:* Experiment 1; *Middle:* Experiment 2; *Right:* Experiment 3. Trial durations are indicated by the thick black lines below the x-axis, and curves extend 2 s before the onset and 12 s after the offset each trial. *Green:* predictions of the 2 temporal-channel model; *Blue:* predictions of the standard model. Both models are fit using data concatenated across Experiments 1 and 2 and then validated against data from Experiment 3.

**Supplementary Figure S2:**
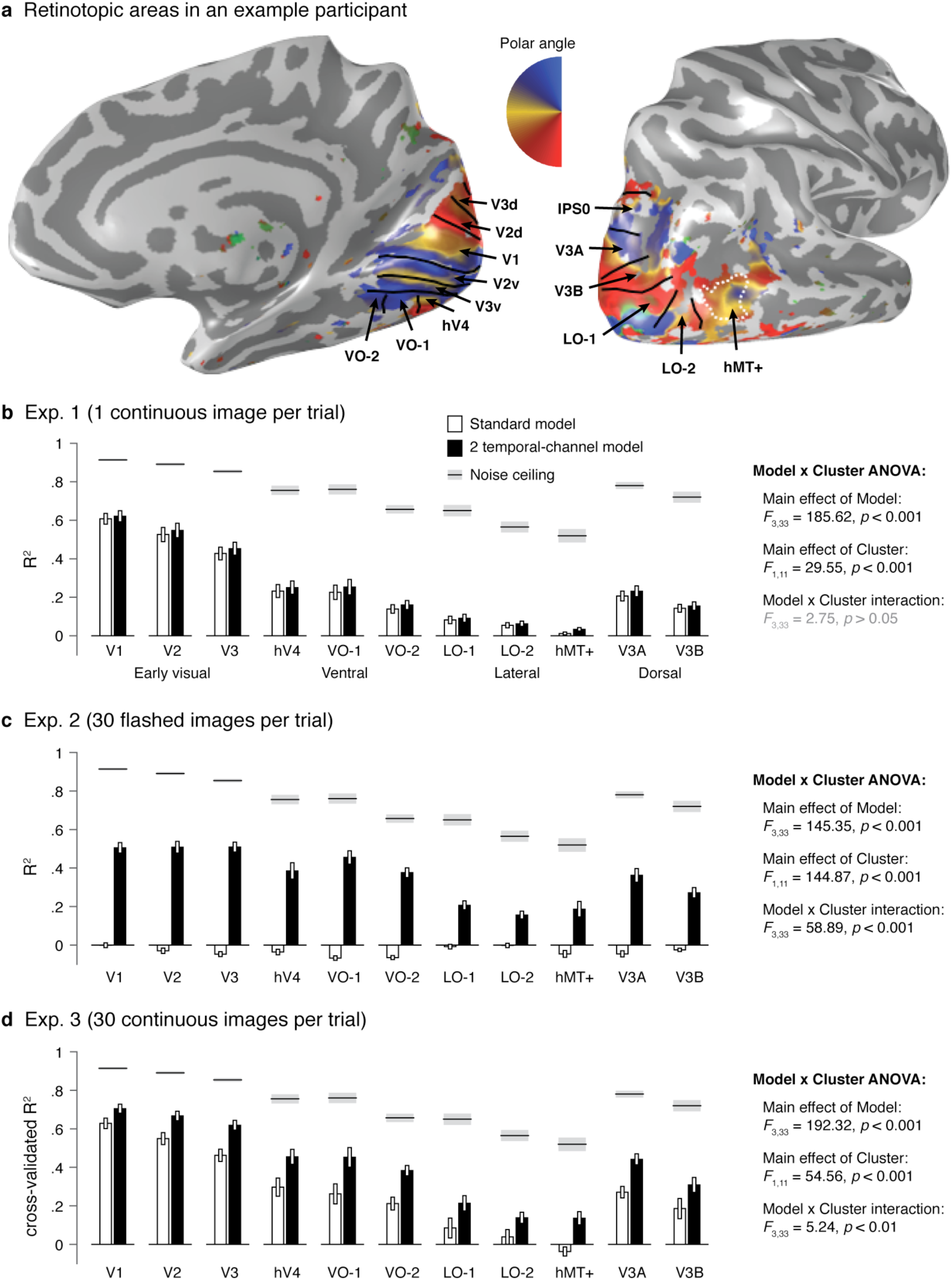
Comparison of the variance explained by the 2 temporal-channel vs. standard model. (a) Definition of retinotopic visual areas in a representative participant. Boundaries of retinotopic areas (*black*) are overlaid on an example participant’s inflated cortical surface depicting a polar angle map generated using pRF mapping. hMT_+_ is outlined with a dashed white line. (**b–d**) The performance of the 2 temporal-channel model and standard model is compared across four clusters of visual areas: *Early visual* (V1, V2, V3), *ventral* (hV4, VO-1, VO-2), *lateral* (LO-1, LO-2, hMT_+_), and *dorsal* (V3A, V3B). Both models are fit separately in each region using data concatenated across Experiments 1 and 2, and the accuracy (*R^2^*) with which these *β* weights predict fMRI responses is quantified separately for (**b**) Experiment 1 and (**c**) Experiment 2. (**d**) Cross-validated *R^2^* of model predictions for independent data in Experiment 3. Note *R^2^* may be negative if the variance in the model prediction exceeds that of the data. *Error bars:* ± 1 SEM across 12 participants. *Black:* 2 temporal-channel model; *White:* standard model; *Gray line:* noise ceiling. Statistical significance of model fit comparison across visual areas was evaluated by a repeated-measures ANOVA for each experiment, shown beside each panel.

**Supplementary Figure S3:**
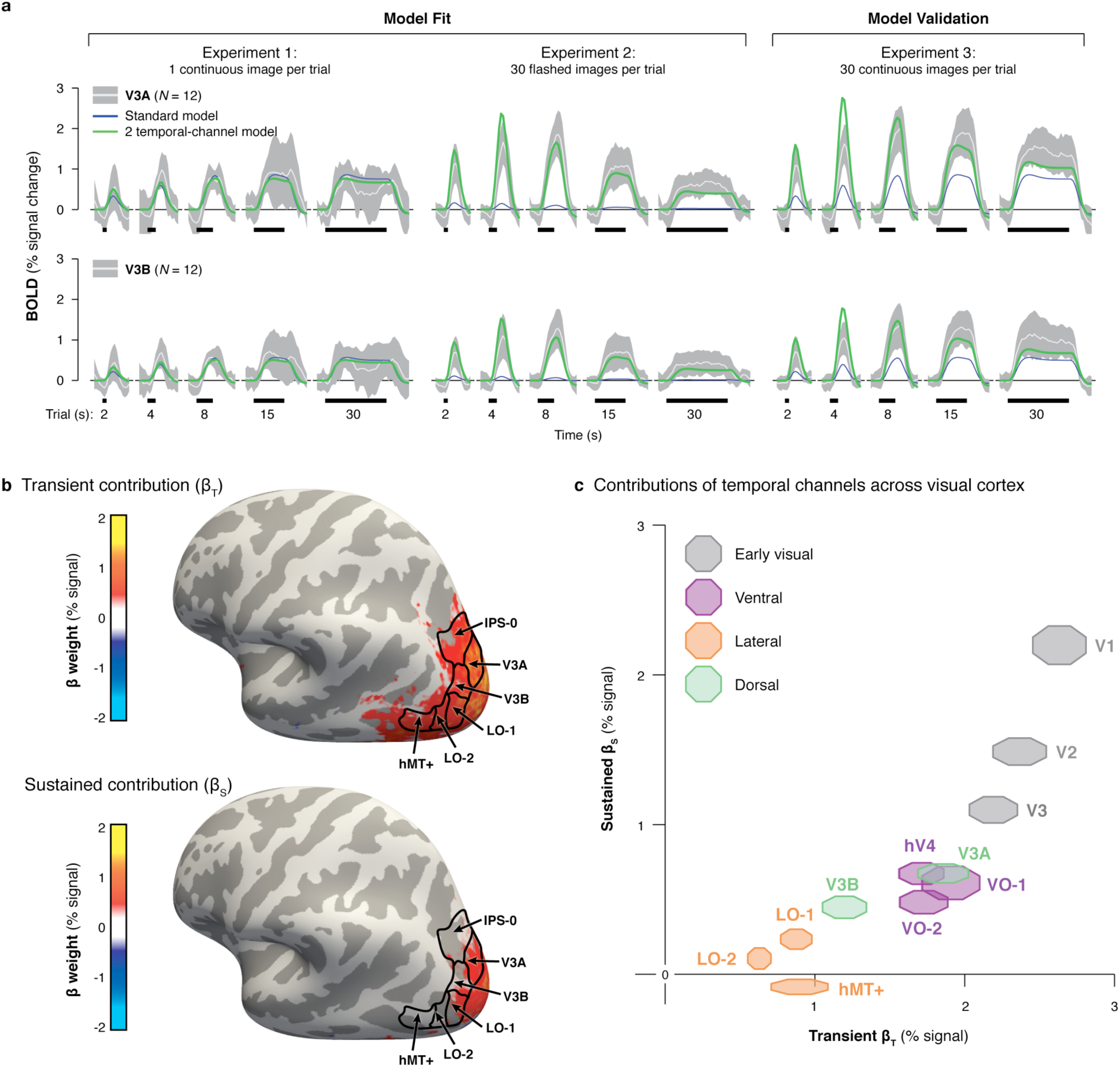
Modeling sustained and transient contributions in dorsal retinotopic visual areas. (**a**) Each row depicts the average ROI response (*white*) ± 1 standard deviation (*gray*) across 12 participants for dorsal regions V3A and V3B. *Left:* Experiment 1; *Middle:* Experiment 2; *Right:* Experiment 3. Trial durations are indicated below the x-axis, and curves extend 2 s before the onset and 12 s after the offset each trial. *Green:* predictions of the 2 temporal-channel model; *Blue:* predictions of the standard model. Both models are fit using data concatenated across Experiments 1 and 2 and then validated against data from Experiment 3. (**b**) Group-averaged (*N* = 12) maps of the transient (*left*) and sustained (*right*) channels shown from a dorsolateral view. We first estimate *β* weights of each channel in each voxel in each participant’s native brain space. *β* weight maps were transformed to the FreeSurfer average brain using cortex-based alignment and averaged across participants in this common cortical space. The resulting group maps were thresholded to exclude voxels with weak contributions (–0.1 > *β* > 0.1). Boundaries of dorsal and lateral regions (*black*) are derived from the Wang Atlas, with hMT_+_ as the union of TO-1 and TO-2. (**c**) The contributions of the transient (x-axis) and sustained (y-axis) channels to responses in each visual area as estimated by the 2 temporal-channel model. Marker size spans ± 1 SEM across 12 participants in each dimension and *β* weights were solved by fitting the 2 temporal-channel using data concatenated across Experiments 1 and 2.

**Supplementary Figure S4:**
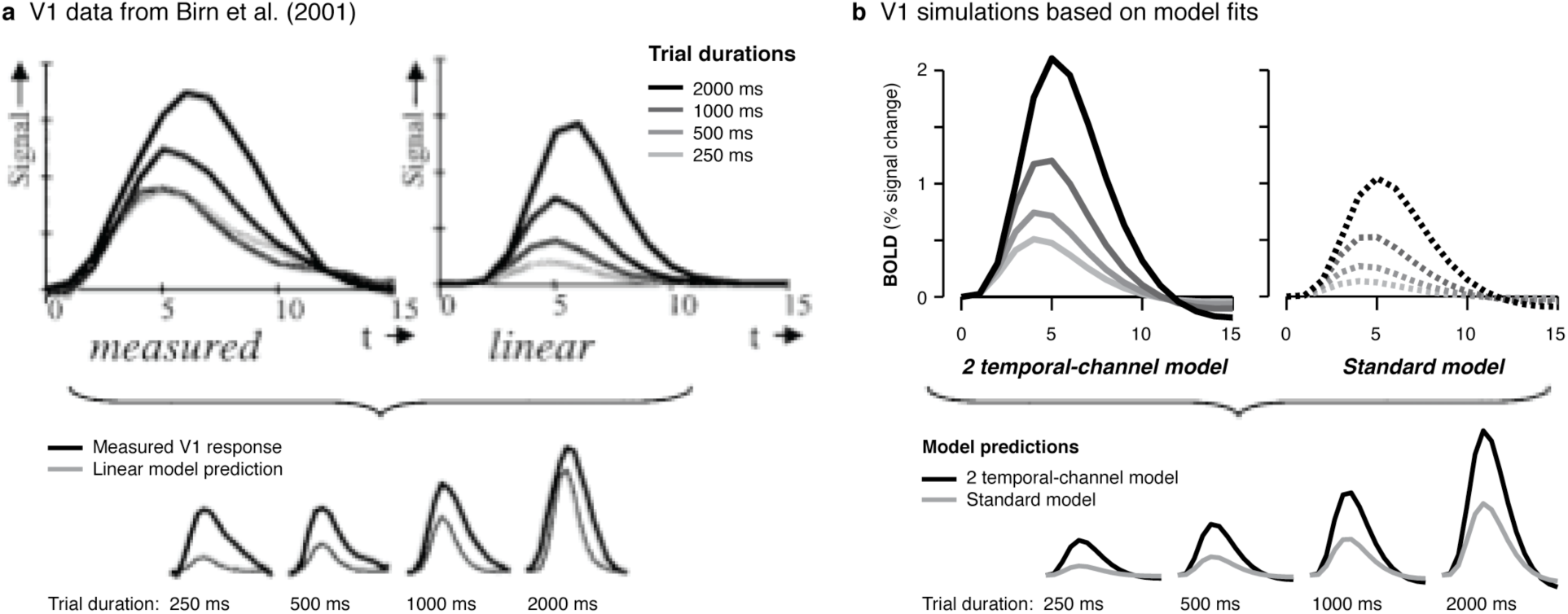
The 2 temporal-channel model explains response nonlinearities for briefly presented stimuli. (**a**)Figure adapted from Birn et al. (y-axis values are unreported in original version). *Top left*: measured V1 responses to brief (250–2000 ms) presentations of a checkerboard stimulus that was contrast inverted at 8 Hz in all trial durations; *Top right:* predicted V1 responses based on a standard linear model solved using responses to longer presentations of the checkerboard stimulus. *Bottom:* same data as above except the measured and predicted fMRI responses are superimposed for each trial duration. (**b**) Simulated V1 responses to the stimuli used by Birn et al. that are derived with the *β* weights solved using models fit to V1 data from Experiments 1 and 2 of the present study. *Left:* predictions of the 2 temporal-channel model for each trial duration; *Right:* predictions of the standard model for each trial duration. *Bottom*: same data as above except the predictions of the two models are superimposed for each trial duration. The simulations show that the standard model replicates Birn et al.’s linear model and underestimate responses. In contrast, the 2 temporal-channel model better explains the measured responses (a-left) and predicts higher responses than the standard model in each duration (b-bottom).

